# Activation of phospholipase C β by Gβγ and Gα_q_ involves C-terminal rearrangement to release auto-inhibition

**DOI:** 10.1101/810994

**Authors:** Isaac Fisher, Meredith Jenkins, Greg Tall, John E Burke, Alan V. Smrcka

## Abstract

Phospholipase C (PLC) enzymes hydrolyse phosphoinositide lipids to inositol phosphates and diacylglycerol. Direct activation of PLCβ by Gα_q_ and/or Gβγ subunits mediates signalling by Gq and some Gi coupled G protein-coupled receptors (GPCRs), respectively. PLCβ isoforms contain a unique C-terminal extension, consisting of proximal and distal C-terminal domains (CTD) separated by a flexible linker. The structure of PLCβ3 bound to Gα_q_ is known, however, for both Gα_q_ and Gβγ, the mechanism for PLCβ activation on membranes is unknown. We examined PLCβ2 dynamics on membranes using hydrogen deuterium exchange mass spectrometry (HDX-MS). Gβγ caused a robust increase in dynamics of the distal C-terminal domain (CTD). Gα_q_ showed decreased deuterium incorporation at the Gα_q_ binding site on PLCβ. *In vitro* Gβγ-dependent activation of PLC is inhibited by the distal CTD. The results suggest that disruption of auto-inhibitory interactions with the CTD, respectively, leads to increased PLCβ hydrolase activity.

## Introduction

Receptor-dependent activation of phospholipase C (PLC) enzymes stimulates hydrolysis of the lipid substrate phosphatidylinositol 4,5 bisphosphate to the second messengers, diacylglycerol (DAG), and inositol 1,4,5 trisphosphate (IP_3_), resulting in downstream activation of protein kinase C (PKC), and calcium release, respectively (Berridge, 1987; Kadamur and Ross, 2013). There are 6 families of PLC isoforms: β, δ, γ, ε, ζ and η (Harden et al., 2011; Kadamur and Ross, 2013; Smrcka et al., 2012). All share a common core domain structure consisting of four EF hand repeats, a TIM barrel catalytic domain, separated into X and Y domains connected by a flexible “X/Y” insert linker, and a C2 domain (Kadamur and Ross, 2013). Most PLC isozymes have a pleckstrin homology (PH) domain at the amino terminus. The individual PLC isotypes differ in their presence or absence of other regulatory domains.

Members of the PLCβ family are among the most well studied PLC isoforms. They are important for a myriad of biological functions including, inflammation (Li et al., 2000), opioid analgesia (Bianchi et al., 2009; Xie et al., 1999), and cardiovascular function (Filtz et al., 2009; Howes et al., 2003; Woodcock et al., 2009; Woodcock et al., 1995) downstream of G protein-coupled receptor (GPCR) activation. There are 4 isoforms of the PLCβ family; 1,2,3 and 4 that are all directly activated by binding Gα_q_ subunits (Lee et al., 1994; Singer et al., 2002; Smrcka et al., 1991; Smrcka and Sternweis, 1993; Taylor et al., 1991). PLCβ2 and β3 are also activated by Gβγ (Lee et al., 1994; Smrcka and Sternweis, 1993). A unique feature of the PLCβ family is a ∼400 amino acid C terminal extension comprised of a well conserved proximal C-terminal domain helix-loop-helix domain, a flexible linker region, followed by a less well conserved distal C-terminal domain (CTD) (Figure 1). The distal CTD has Bin, Amphiphysin and Rvs. (BAR) domain homology and folds into a helical bundle (Lyon et al., 2013; Singer et al., 2002). The first structure of PLCβ3 bound to Gα_q_ shows direct interactions of Gα_q_ with the proximal CTD and the EF hands (Waldo et al., 2010). A more recent structure of full length PLCβ3 includes the extended C-terminus and can serve as a model for the domain organization of all PLCβ isoforms (Fig. 1) (Lyon et al., 2011). Both the proximal and distal CTD are required for activation by Gα_q_, however the distal CTD is dispensable for Gβγ activation (Lee et al., 1993; Wu et al., 1993).

**Figure 1:**
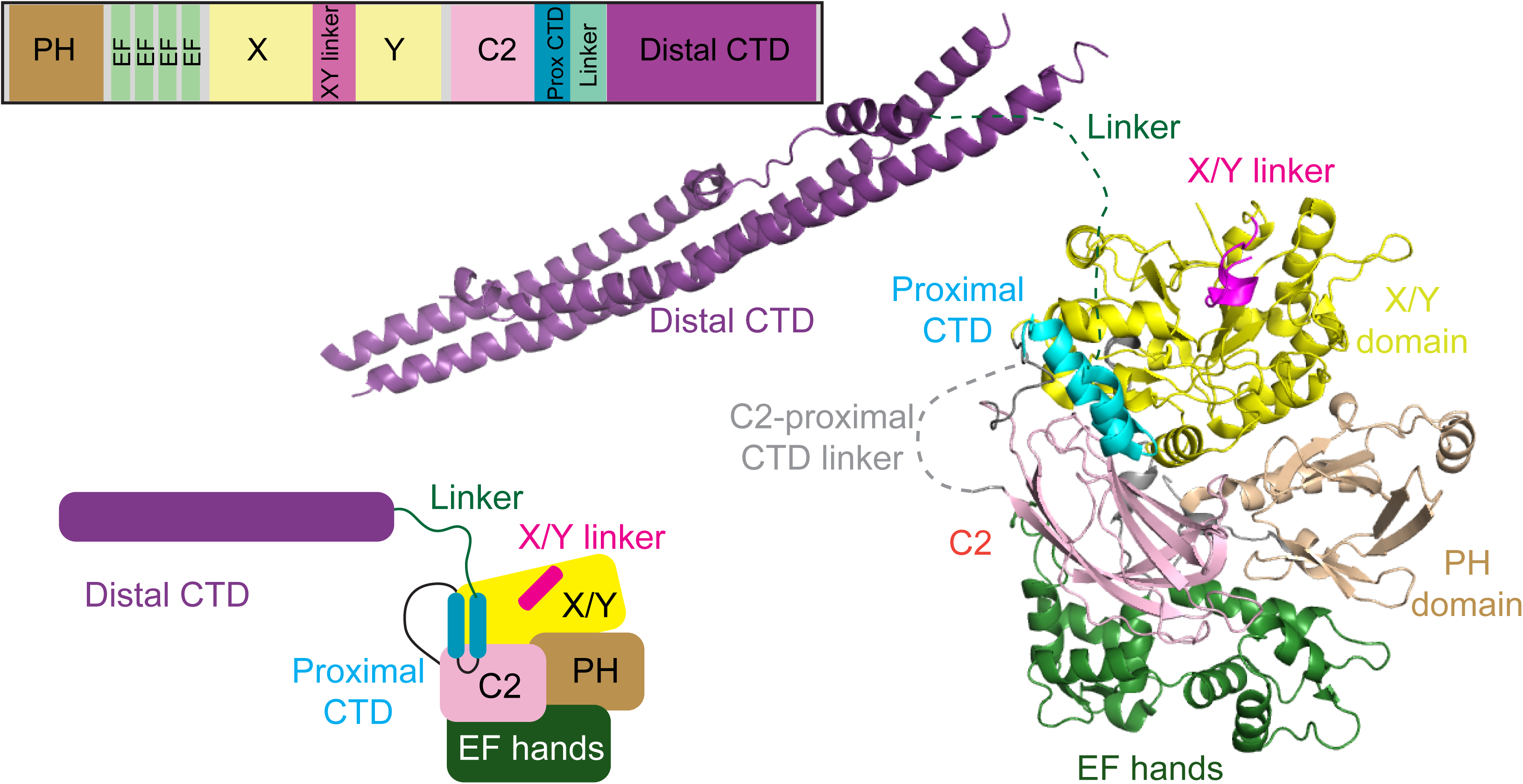
Domain Architecture and model of PLCβ2 architecture. A model of PLCβ2 based on a monomer of PLCβ3 from the co-crystal structure of Gα_q_-PLCβ3 (PDB:4GNK), and the proximal CTD from the structure of *S. officinalis* PLC21 (PDB: 3QR0). This model is oriented such that the active site is in close proximity to the plasma membrane. Domains are colored as in the domain schematics in the top and bottom left. A model of the structure is shown on the right, with this used as a template for describing all HDX-MS data.

There are two proposed general mechanisms of PLCβ activation, with multiple models proposed for how GPCRs mediate this activation. The binding site for Gα_q_ bound to PLCβ is well defined, but no clear binding site has been identified for Gβγ. Proposed mechanisms of activation involve either the X/Y linker or the proximal CTD (also referred to as the helix loop helix region). The negatively charged X/Y linker acts as a lid on the active site occluding access to PIP_2_ substrate. It has been proposed that negative charges on the flexible region of the linker are repelled upon membrane binding, or reorientation on the membrane surface, and this acts to remove the “lid” allowing for substrate (PIP_2_) to bind the catalytic site (Hicks et al., 2008). Genetic deletion of the flexible X/Y linker leads to an increase in basal activity, but this activity can be further stimulated by G-proteins, suggesting there is more to G-protein regulation of PLC activity than removal of the X/Y linker steric occlusion of the active site. A second proposed mechanism for activation involves the proximal CTD being autoinhibitory when packed against the catalytic core of PLC. In an inactive structure of invertebrate PLC (PLC21) (PDB: 3QR0 and 3QR1) the proximal CTD interacts with the core of the enzyme (Lyon et al., 2013; Lyon et al., 2011). In Gα_q_-PLCβ3 co-crystal structures the proximal CTD is pulled away from the core via binding to Gα, leading to disruption of the autoinhibitory interface (Lyon et al., 2011; Waldo et al., 2010). Supporting this hypothesis, mutations that interfere with the proximal CTD-core interactions cause a robust increase in activity that is not further stimulated by Gα_q_.

There are two proposed binding sites of Gβγ on PLCβ. One of the proposed binding sites for Gβγ on PLCβ is in the PH domain. Indeed, isolated PH domain has been shown to inhibit Gβγ stimulated PLC activity by competing for binding to Gβγ and chimeras containing the PH domain of PLCβ2 on PLCδ can confer the ability to be activated by Gβγ subunits (Han et al., 2011; Kadamur and Ross, 2016; Wang et al., 2000). Interestingly, one proposed PH domain binding site is not surface exposed in structural models of PLC, with binding of Gβγ proposed to be mediated by dynamic dissociation from the PLC core, leading to exposure of the Gβγ-binding surface on the PH domain (Kadamur and Ross, 2016). Cross-linking to restrict this putative PH-domain movement inhibits both Gβy binding to, and activation of, PLCβ. The other proposed binding site of Gβγ on PLCβ is in the Y-domain. Mutational analysis of this region decreases Gβγ activation (Bonacci et al., 2005), and peptides corresponding to this region inhibits Gβγ-dependent activation of PLCβ2, and can be directly crosslinked to Gβγ (Kuang et al., 1996; Sankaran et al., 1998). Complicating analysis of the molecular basis for G protein-dependent activation of PLC, is that the activation occurs on a membrane surface, and studying these interactions by standard biophysical methods is extremely challenging.

To further define the molecular mechanisms that mediate activation of PLCβ2 by GPCRs on membranes we utilized hydrogen deuterium exchange mass spectrometry (HDX-MS). This technique has provided key insight into the molecular mechanisms of activation of lipid signalling enzymes downstream of GPCRs (Dbouk et al., 2012; Vadas et al., 2013). Our analysis reveals unique insights into the role of C-terminal domains of PLCβ2 in activation by both Gα_q_ and Gβγ. The results show that Gβγ causes large scale allosteric rearrangement of the distal CTD domain, and that Gα_q_ interacts with the distal CTD as well as the proximal CTD. Biochemical assays reveal a novel direct CTD-catalytic core interaction that inhibits PLC activation. Overall, this work provides fundamental novel insight into unique regulatory mechanisms that control PLCβ activation by GPCRs, and indicates a unifying role for the C-terminal extension in mediating G-protein activation.

## Results

Activation of PLC by G proteins requires a membrane surface (Charpentier et al., 2014). High resolution structural studies of peripheral membrane proteins are generally not compatible with, or are very difficult to perform on membrane surfaces. We used hydrogen-deuterium exchange mass spectrometry (HDX-MS) to examine the dynamics of PLCβ2 in the presence and absence of membranes, Gβγ, and activated Gα_q_. HDX-MS measures the exchange rate of amide hydrogens with deuterated solvent, and as the main determinant of exchange is the stability of secondary structure, it is an excellent probe of conformational dynamics. Deuterium incorporation is localised at peptide level resolution through the generation of pepsin generated peptides. HDX-MS coverage of PLCβ2 was excellent, with 232 identified peptides that spanned 92% of the primary sequence of PLCβ2. HDX-MS experiments compared the dynamics of PLCβ2 on and off lipid vesicles composed of phosphatidylethanolamine(PE), phosphatidylserine (PS), and phosphatidylinositol 4,5 bisphosphate (PIP_2_), and in the presence or absence of either Gβγ or activated Gα_q_.

### HDX-MS reveals conformational changes in the C-terminus and PLCβ core that occur upon membrane binding

Coincubation of PLCβ2 with PS/PE/PIP_2_ containing lipid vesicles revealed multiple regions of significantly decreased deuterium exchange (defined as changes greater than both 4% and 0.4 Da at any timepoint with a student t-test value less than p<0.05) that occurred across multiple domains of PLCβ2 (Figure 2A-C). There were extensive decreases in exchange that occurred on all helices of the distal CTD, spanning regions 878-931, 1020-1109 and 1169-1185. This is consistent with direct interactions of lipid with the distal CTD, which has been shown to be a primary determinant of membrane binding in PLCβ (Adjobo-Hermans et al., 2013; Kim et al., 1996; Wu et al., 1993). It is possible that some of these changes also may represent conformational changes mediated by interactions of the distal CTD with the PLC core that occur upon membrane binding, since changes were seen on multiple sides of the CTD domain (Lyon et al., 2013). There were also decreases in exchange in regions spanning the PLCβ2 core, including in the EF hand domain (residues 255-290) and C2 domain (694-701). These changes were surprising since these regions would not be predicted to participate in membrane binding based on previous crystallographic studies. Intriguingly, both of these regions are very close to the C-terminus of the C2 domain, which is immediately followed by the proximal CTD domain (Figure 1, Figure 2A). This suggests that these regions are involved in allosteric changes in PLC associated with membrane binding. The region in the C2 domain with decreases in exchange would be predicted to be directly in contact with the putative autoinhibitory proximal CTD binding site based on the PLC21 structure (Lyon et al., 2011), suggesting a possible change in the interaction of the proximal CTD with the catalytic core upon membrane binding.

**Figure 2.**
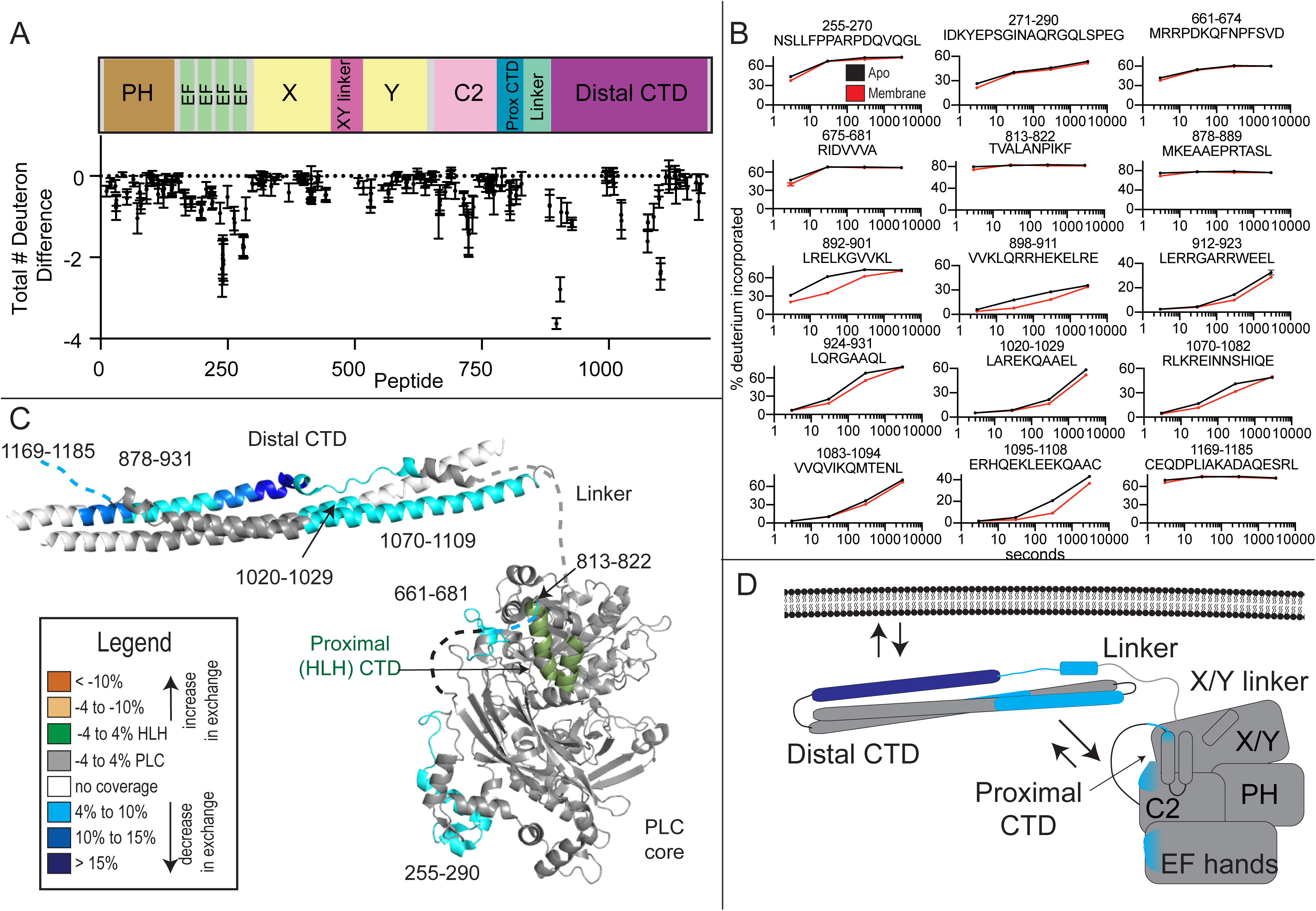
Membrane dependent changes in PLCβ2 dynamics. **(A)** The total number of deuteron difference between apo and membrane bound states for all peptides analyzed over the entire deuterium exchange time course for in PLCβ2. Every point represents the centroid of an individual peptide, with the domain schematic present above. Error bars represent S.D. (n=3). **(B)** Representative PLCβ2 peptides displaying increases or decreases in exchange in the presence of membrane are shown. For all panels, error bars show SD (n=3), with most smaller than the size of the point. The full list of all peptides and their deuterium incorporation is shown in the source data. **(C)** Peptides with significant changes in deuterium incorporation (both >0.4 Da and >4% and T-test p < 0.05 at time point) in the presence of membrane are mapped on a structural model of PLCβ2 as described in Fig. 1. Differences are mapped according to the legend. **(D)** Cartoon schematic displaying deuterium exchange differences, and potential mechanisms for altered protein dynamics.

We saw no changes in deuterium exchange consistent with the X/Y linker displacement by the membrane surface, but exchange rates in this area were extremely rapid indicating a high degree of flexibility of this region of the protein. We observed decreased deuterium incorporation in the proximal CTD and its linker to the C2 domain of PLCβ2 (residues 813-822). This may reflect an altered conformation of this region upon binding to membrane or direct interaction with membranes. The helix-loop-helix proximal CTD directly interacts with Gα_q_, and it is possible that this altered conformation may prime the proximal CTD for interaction with Gα_q_.

### Conformational changes that result from Gα_q_ binding to PLCβ on membranes

Next, we compared HDX rate differences between PLC-liposome complexes to PLC-Gα_q_-liposome complexes. There were extensive decreases in exchange at regions that correspond directly to the known binding site of Gα_q_ on PLC as determined by X-ray crystallography (Lyon et al., 2013; Lyon et al., 2011), strongly supporting the validity of this method and our approach. This included the proximal CTD, which is the principle binding site for Gα_q_ (Lyon et al., 2013; Waldo et al., 2010) showing a >40% decrease in deuterium incorporation (Figure 3A-D). In addition, regions of the EF hand (residues 255-272), and C2 domain (675-681, 740-745, and 802-812), either at the interface with Gα_q_, or directly adjacent, also showed significant decreases in exchange upon binding to Gα_q_ on membranes. This region of the EF hand directly corresponds to structural elements directly involved activation of the GTPase activity of Gα_q_ (Waldo et al., 2010). There was also an increase in exchange in a region of the EF hands (271-290) adjacent to the Gα_q_ binding interface, revealing allosteric conformational changes that occur upon Gα_q_ binding. The distal CTD showed decreases in exchange directly at the contact site with the Gα_q_ N-terminal region (CTD region 1070-1109), as well as other regions directly adjacent (878-889 and 1020-1029). Gα_q_ appears to contact the N-terminal helix in the Gα_q_-full length PLCβ3 co-crystal structure but there was some concern that this may represent a crystal packing artefact (Lyon et al., 2013). The results from the HDX experiments confirm that in solution, in the presence of membranes, many of the contacts seen in the X-ray crystal structures are valid contact sites. Additional conformational alterations are revealed that may be critical for activation.

**Figure 3.**
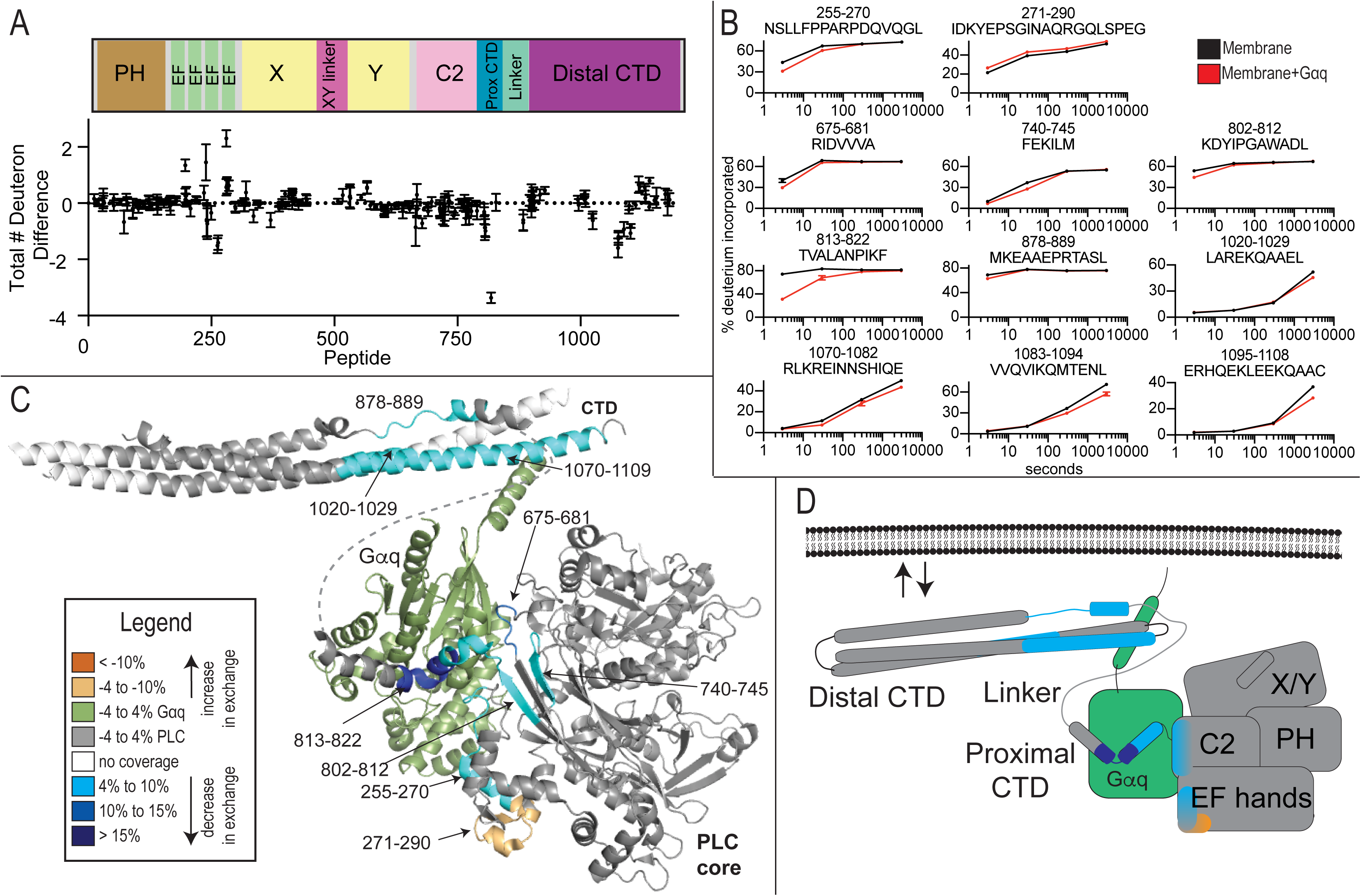
Gα_q_ dependent changes in PLCβ2 dynamics. **(A)** The total number of deuteron difference between membrane and membrane*-Gα_q_* states for all peptides analyzed over the entire deuterium exchange time course for PLCβ2. Every point represents the centroid of an individual peptide. Error bars represent S.D. (n=3). **(B)** Representative PLCβ2 peptides displaying increases or decreases in exchange in the presence of *G*α_q_ are shown. For all peptides, error bars show SD (n=3), with most smaller than the size of the point. The full list of all peptides and their deuterium incorporation is shown in the source data. **(C)** Peptides with significant changes in deuterium incorporation (both >0.4 Da and >4% and T-test p < 0.05 at time point) in the presence of *G*α_q_ are mapped on a structural model of PLCβ2 as described in Fig. 1. Differences are mapped according to the legend. **(D)** Cartoon schematic displaying deuterium exchange differences on a model of *G*α_q_ binding.

### Gβγ binding to PLCβ on membranes reveals extensive conformational changes in the distal CTD

Next, we compared changes in deuterium incorporation between PLC-membrane complexes and PLC-Gβγ-membrane complexes. The most striking and unexpected effect of Gβγ binding to PLCβ was increased deuterium exchange on almost the entire distal CTD of PLCβ2 (multiple peptides spanning 892-1114) (Figure 4A-D). This is not due to competition for membrane binding because the other membrane associated changes in dynamics i.e. EF hand and C2 domain do not show this increase. (Figure 4A-D). Additionally, even under the high protein concentrations used for HDX experiments, Gβγ still increased PLC activity 3-4-fold (Supplemental figure 1). This indicates that Gβγ causes PLCβ2 to undergo a conformational change that rearranges the distal CTD. One possibility is that Gβγ binding leads to disruption of an auto-inhibitory interaction of the CTD with the PLCβ2 core. No changes were seen in the PLCβ2 core that would be consistent with this model. However; disruption of intramolecular interactions of the CTD with the core, with concomitant binding of Gβγ to the core would result in no net change in deuterium incorporation in the core.

**Figure 4.**
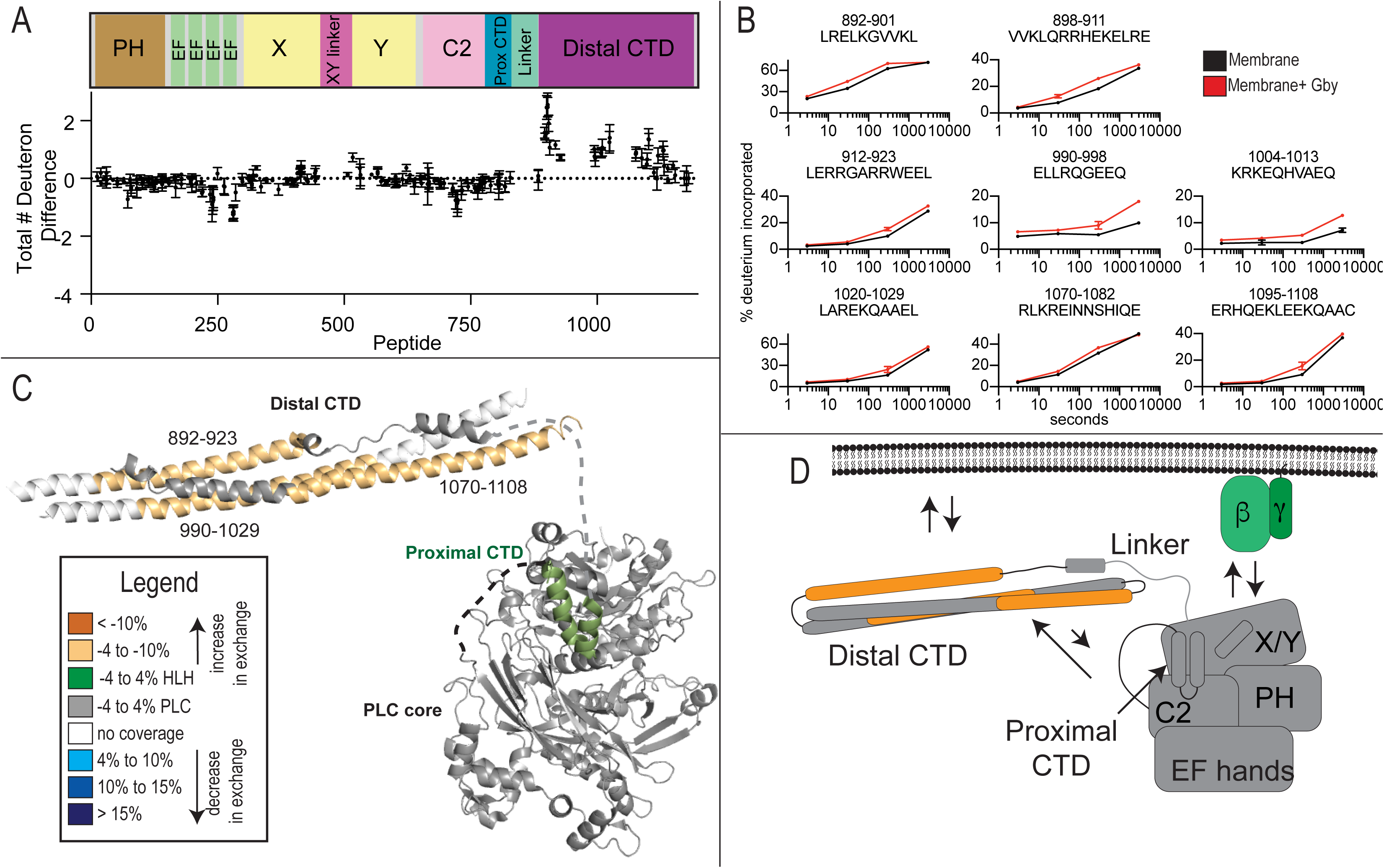
Gβγ binding to PLCβ2 causes conformational changes in the distal CTD domain. **(A)**The total number of deuteron difference between membrane and membrane*-* Gβγ states for all peptides analyzed over the entire deuterium exchange time course for PLCβ2. Every point represents the centroid of an individual peptide. Error bars represent S.D. (n=3). **(B)** Representative PLCβ2 peptides displaying increases or decreases in exchange in the presence of Gβγ are shown. For all panels, error bars show SD (n=3), with most smaller than the size of the point. The full list of all peptides and their deuterium incorporation is shown in the source data. (**C)** Peptides with significant changes in deuterium incorporation (both >0.4 Da and >4% T-test p < 0.05 at time point) in the presence of membrane are mapped on a structural model of PLCβ2 as described in Fig. 1. Differences are mapped according to the legend. **(D)** Cartoon schematic displaying deuterium exchange differences upon Gβγ, and potential mechanisms for altered protein dynamics.

### The distal CTD is autoinhibitory

To test the hypothesis that the distal CTD was autoinhibitory, we generated a PLCβ2 mutation deleting the distal CTD (PLCβ2-ΔCTD). A diagram of this truncation is shown in Figure 5A. We then examined the ability of PLCβ2-ΔCTD to be activated by Gβγ. We incubated varying concentrations of Gβγ with PLCβ2-ΔCTD or full-length PLCβ2 in a PLC enzyme activity assay using phospholipid vesicles containing PIP_2_ substrate. We predicted that the distal CTD-deleted PLC would have increased responsiveness to Gβγ stimulation as it would lack one of the autoinhibitory constraints opposing activation by Gβγ. Gβγ activated PLCβ2-ΔCTD with increased efficacy but no change in the EC50 (Figure 5B). Basal Ca^2+^ stimulated activity of PLCβ2-ΔCTD was similar to that of full-length PLCβ2 (supplemental figure 2). These data suggest that the distal CTD may act an autoinhibitory domain.

**Figure 5:**
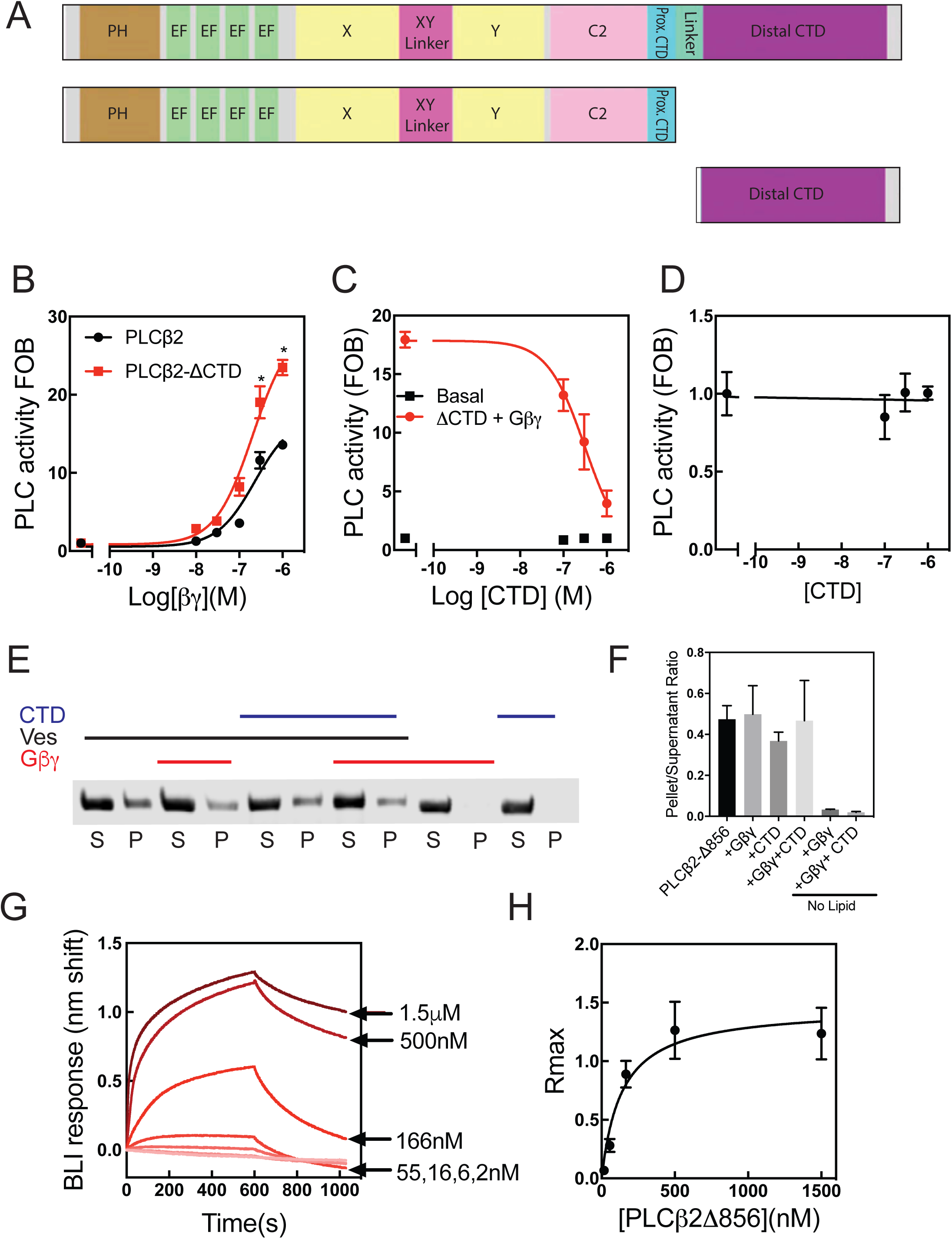
The Distal CTD autoinhibits Gβγ activation of PLCβ2. **(A)** Schematic of PLCβ2 constructs. **(B)** Assay of PLC enzymatic activity for the indicated purified PLC protein with the indicated concentrations of purified Gβγ subunits performed at least 3 times in duplicate. The data shown are mean ± S.E. and analysed by 2-way ANOVA with multiple comparisons post test. *, p < 0.05 versus Full length PLCβ2 at equal concentration of Gβγ. **(C)** Assay of PLC enzymatic activity for the PLCβ-Δ856 with 250 nM Gβγ subunits and incubated with the indicated concentrations of purified CTD performed at least 3 times in duplicate. **(D)** Reconstituted assay of PLC enzymatic activity for the PLC-Δ856 basal activity incubated with the indicated concentrations of purified CTD performed at least 3 times in duplicate. (Basal CPM = 234±15, total CPM = 6700 in the assay). **(E)** Representative western blot image of PLCβ following ultracentrifugation in the presence (+) or absence (−) of PE/PIP_2_ liposomes, 1 μM CTD and/or purified 500ng Gβγ as indicated. **(F)** The ratio of protein present in the supernatant samples versus the respective total protein control samples is shown. No significant differences in lipid binding for PLCβ2 in the presence of CTD or Gβγ was detected. Data represent at least three independent experiments. **(G)** Representative sensorgram of PLCβ2-Δ856 binding to purified CTD via Biolayer interferometry (BLI). **(H)** Binding isotherm of maximal PLCβ2-Δ856 binding to CTD at indicated concentrations. Data represents at least three independent experiments.

To provide further evidence that the distal CTD is autoinhibitory, we tested whether the addition of purified distal CTD could inhibit Gβγ stimulated PLCβ activity when incubated with PLCβ2-ΔCTD in trans. We expressed and purified the triple helical bundle of the distal CTD (residues 864-1184) and incubated varying concentrations of purified CTD with Gβγ and PLCβ2-ΔCTD and measured PLC enzyme activity. We predicted that the distal CTD would inhibit Gβγ-dependent PLC activation. The purified CTD inhibited Gβγ stimulated activity with an IC50 of 194 ±46 nM while having no appreciable effect on basal activity (Figure 5C and D).

One possible explanation of these results is that, since the CTD binds lipids, the CTD added in trans could compete for binding of PLC to the vesicle surface and thereby inhibit PLC activity. The observation that CTD addition did not affect PLC basal activity argues against this hypothesis, but to further test this idea, we measured binding of purified PLCβ2 binding to lipid vesicles, in the presence or absence of purified CTD, using a centrifugation-based lipid binding assay. At the highest concentration of CTD tested in the PLC activity reconstitution assay, the CTD had no significant effect on lipid binding of PLCβ2-ΔCTD (Figure5 E, F).

In order to determine if the CTD of PLC directly interacts with the catalytic core of PLCβ, we used biolayer interferometry (BLI) to examine direct binding. We immobilized the CTD of PLCβ2 and examined binding to increasing concentrations of purified PLCβ2-ΔCTD. PLCβ2-ΔCTD bound directly to the CTD of PLCβ2 with a Kd of 146±88 nM. The response reached equilibrium and exhibited little non-specific binding to the reference tip indicating this is a specific interaction (Figure 5F, G). Taken together, these data suggest the CTD is inhibitory to PLC activity, and this inhibition is through direct binding of the distal CTD to the catalytic core of PLCβ2

### CTD movement is important for activation of PLCβ

To determine if the movement of the distal CTD upon Gβγ binding observed by HDX-MS is important for activation we generated a construct lacking the flexible linker between the proximal CTD and distal CTD (PLCβ2-Δ-linker) (Figure 6A). We hypothesized that deleting this linker would prevent the distal CTD from forming auto-inhibitory interactions with the catalytic core due to decreased conformational flexibility (Figure 6B). If movement of the distal CTD was important for activation, it would be expected that PLCβ2-Δ-linker would have increased basal activity, and would not be further activated by Gβγ. We examined activation of PLCβ2-Δ-linker by G-proteins in a cos-7 cell co-transfection assay. As expected, PLCβ2-Δ-linker had increased basal activity that was not increased in the presence of either Gβγ or Gα_q_ (Figure 6C). Since no further increase was seen in activity in the presence of either G-protein, this suggests a unified mechanism where the orientation of the proximal and distal CTD domains is critical for PLCβ2 activation by heterotrimeric G proteins. These data also agree with the changes in dynamics we see in HDX with Gβγ suggesting that the rearrangement of the distal CTD is an integral step in Gβγ activation.

**Figure 6:**
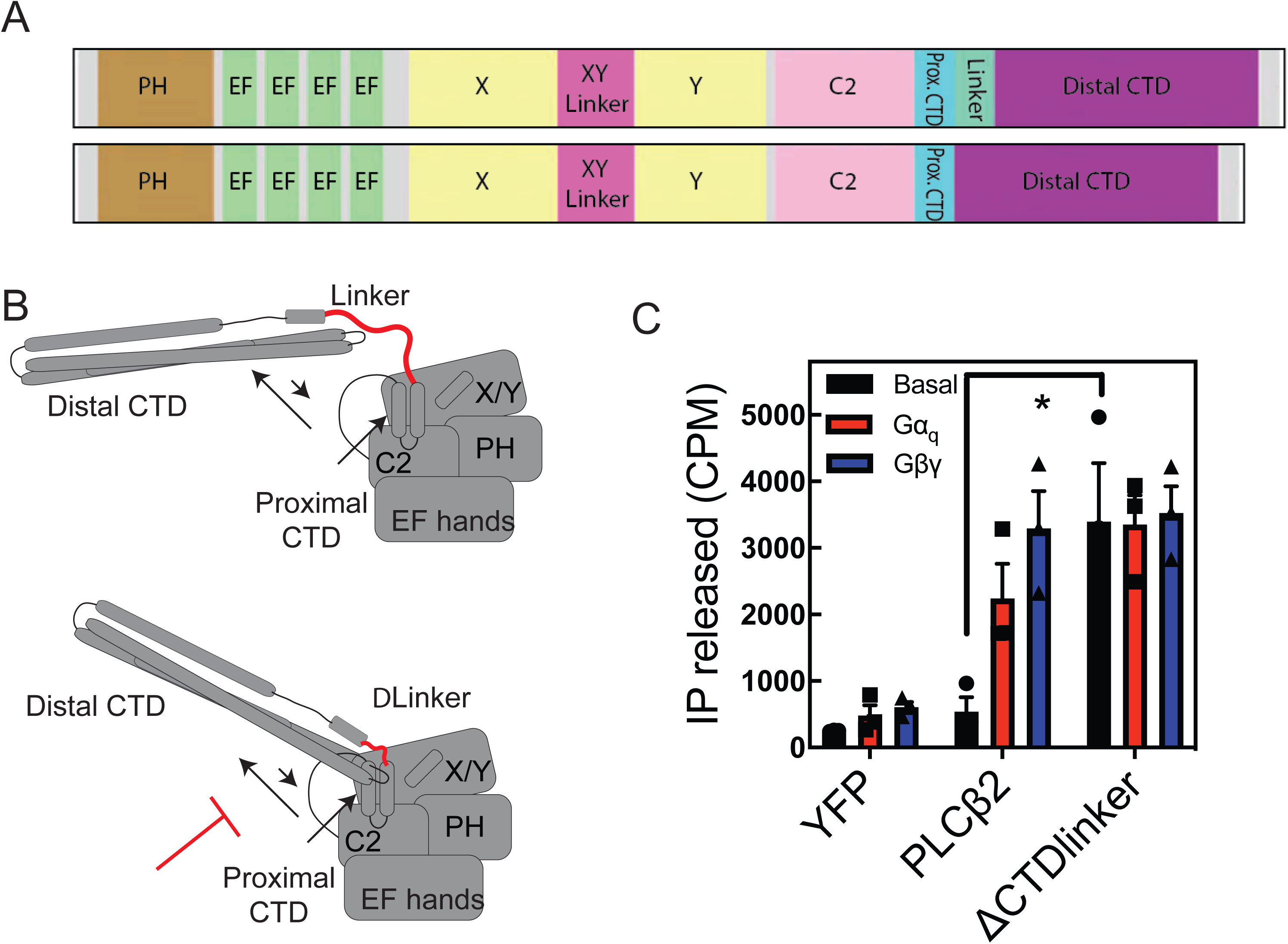
Restriction of Distal CTD movement increases PLC activity. **(A)** Schematic of CTD-linker deletion. **(B)** Cartoon diagram of model of CTD-linker deletion effects. By preventing C-terminal domain rearrangement, CTD linker would prevent autoinhibition. **(C)** Total inositol phosphate assay. COS-7 cells were transfected with 250 ng PLCβ2 WT or PLCβ2-ΔCTD-linker in the presence or absence of 200 ng Gβ_1_ and 200 ng Gγ_2_ or 200ng Gα_q_, and total [^3^H] inositol phosphate accumulation was measured. The data shown are mean ± S.E. for at least three independent experiments and analysed by 2-way ANOVA with multiple comparisons. *, p < 0.05

### Proximal CTD inhibits PLC activation

As the mechanism of activation of PLCβ by Gα_q_ has been established to use both the distal and proximal CTD, and these two domains are covalently tethered, we hypothesized that distal CTD rearrangement could destabilize the proximal CTD, preventing it from adopting an autoinhibited state. We tested PLCβ2 with a c-terminal deletion of both the proximal CTD and distal CTD (PLCβ2-Δ818) and a mutation (PLCβ2-P819A) predicted to disrupt formation of the helix loop helix conformation of the proximal CTD that is stabilized upon Gα_q_ binding (Figure 7A). Both of these mutations greatly reduced basal PLC activity, but also prevented activation by Gα_q_, and, surprisingly, Rac1, seen as a significant reduction in the stimulated activity relative to basal activity when Gα_q_ and Rac1 are cotransfected. (Figure 7B, C). This indicates this region is important for activation of PLCβ2 by these G proteins. While the Gβγ-stimulated activity was reduced (probably as a result of the overall reduction in the PLC activity of these mutants), there was little effect on the ability of Gβγ to increase PLC activity relative to the basal activity of that mutant. This suggests less reliance on proximal CTD interactions with the core for the mechanism for activation by Gβγ.

**Figure 7:**
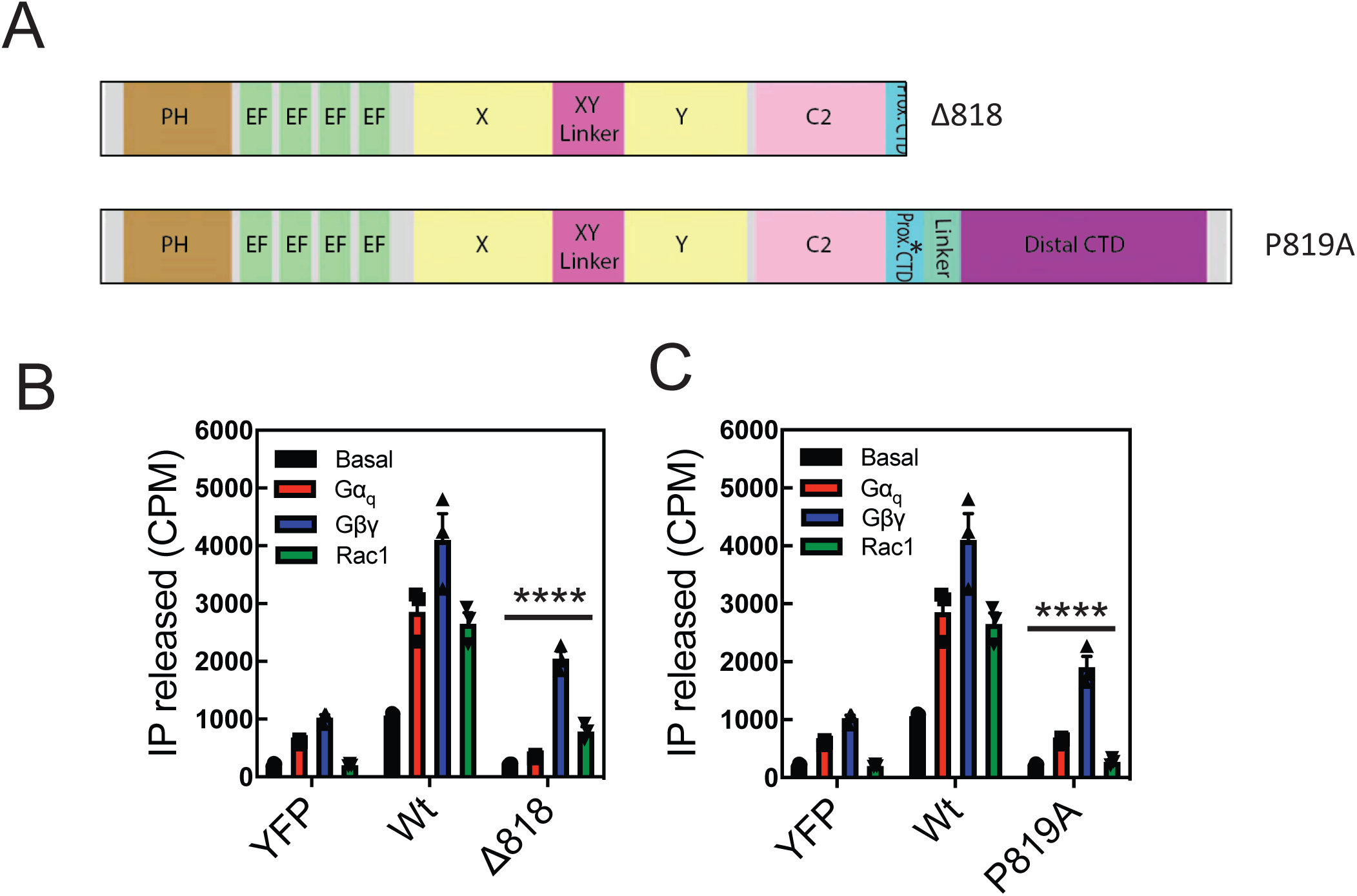
The Proximal CTD is involved PLC activation by Gα_q_ and Rac1. **(A)** Schematic of mutants and deletions of PLCβ2. **(B)** COS-7 cells were transfected with 250ng of WT PLCβ2 P819A PLCβ2 in the presence or absence of 200ng Gβ_1_ and 200ng Gγ_2_ or 300 ng Rac1G12V or 200ng Gα_q_ and total [^3^H] inositol phosphate accumulation was measured. **(C)** COS-7 cells were transfected with 250ng of WT PLCβ2 or Δ818-PLCβ2 in the presence or absence of 200ng Gβ_1_ and 200ng Gγ_2_ or 300 ng Rac1G12V or 200 ng Gα_q_ and total [^3^H] inositol phosphate accumulation was measured. For all panels the data shown are mean ± S.E. for at least three independent experiments and analysed by 2-way ANOVA with multiple comparisons. ****, p < 0.001 statistically different that wt PLC transfection.

## Discussion

Here we used HDX-MS to examine the molecular mechanisms that mediate PLCβ3 activation by GPCR subunits Gα_q_/Gβγ on membranes. Intriguingly, we find that Gβγ binding causes dramatic conformational rearrangement of the distal CTD, suggesting that the CTD is involved in activation by Gβγ. To provide support for this idea we show 1) that the CTD can bind directly to the catalytic core in-trans to inhibit PLC activity, and 2) restricting interactions of the CTD with the catalytic core through removal of the flexible linker strongly increased PLCβ activity that was not further activated by G proteins. Others have shown that deleting the linker of PLCβ3 increases basal activity and this construct cannot be further activated by Gα_q_ (Lyon et al., 2013), but here we demonstrate that PLCβ2 with the CTD-linker deleted did not respond to either heterotrimeric G-protein, suggesting that movement of the distal CTD is important for activation by both G-proteins.

Deletion or mutation of the proximal CTD did not dramatically inhibit Gβγ-dependent activation of PLC, suggesting that rearrangement of the proximal CTD is not required for Gβγ activation. These data indicate that rearrangement of either the proximal or distal CTDs are generally involved in all activation processes, while proximal CTD rearrangement is critical for regulation of activity Gα_q_ and Rac but not Gβγ.

It is interesting to note incubation with phospholipid vesicles caused decreased deuterium exchange in several domains not thought to be involved in membrane binding. Domains far from the active site and the C-terminus that show decreased dynamics on lipid vesicles include the EF hands and the proximal CTD. This suggests that there are conformational changes in PLC that occur upon membrane binding, and this might be due to either changed inter-domain interfaces or conformational changes in PLC upon membrane binding. Since membranes are critical for the allosteric activation of PLCβ enzymes by G-proteins, these conformational changes could be essential for activation by Gβγ or Gα, as predicted for G-protein activation of class I PI3Ks (Dbouk et al., 2012; Vadas et al., 2013). It has been shown that conformational flexibility involving EF hands 3 and 4, and the PH domain are important for activity of multiple PLC variants (Garland-Kuntz et al., 2018).

Surprisingly, we did not observe G-protein or membrane dependent changes in deuterium exchange in either the X/Y linker, PH domain, or the Y domain. There were no changes in the dynamics of the interface between the PH domain and EF hand in the presence of membranes or G proteins. Overall our data does not provide evidence to support rearrangement of the X/Y linker by the membrane, a dynamic PH-EF hand interface for Gβγ binding, or binding of Gβγ to the Y domain, but the HDX-MS data are not incompatible with these processes.

A surprise from our data is that activation of PLCβ by different G-proteins converge on a unifying mechanism involving disruption of inhibition by the c-terminal extension, with the different proximal and distal CTD domains participating in different ways. On membrane surfaces, Gα_q_ interfaces directly with the proximal CTD and distal CTD, with these interactions disrupting all auto-inhibitory interactions. On membranes Gβγ disrupts auto-inhibition by the distal CTD, most likely due to it forming a direct interface with a portion of the PLCβ core. The direct interface of the Gβγ interface with the PLC core is unknown. It is unclear why no regions were protected from exchange by Gβγ binding. It is possible that the region exposed upon disruption of the distal CTD domain inhibitory interface is what is responsible for Gβγ binding, and this would lead to no overall difference in deuterium incorporation.

Our data, taken together, suggest an activation mechanism of PLC by Gβγ where Gβγ causes a conformational rearrangement of the distal CTD through an allosteric mechanism, breaking the interaction between the catalytic core and the distal CTD, while Gα_q_ binds directly to both the proximal and distal CTD in order to break autoinhibitory interactions of the C-terminal domain, in this way activation of PLCβ by either G-protein involves C-terminal domain rearrangement but each G-protein engages this regulatory region of PLC differently, Gα_q_ utilizing direct interaction while Gβγ somehow coordinates a conformational change involving this region. (Figure 8). Further study will be required to conclusively define the exact molecular mechanisms by which Gβγ both directly binds to PLCβ, and releases auto-inhibition.

## Methods

### Transfection of COS-7 cells and quantitation of phospholipase C activity

All cell culture reagents were obtained from Invitrogen. COS-7 cells were obtained from ATCC. COS-7 cells were seeded in 12-well culture dishes at a density of 100,000 cells per well and maintained in high-glucose Dulbecco’s modified Eagle’s medium containing 5% fetal bovine serum, 100 units/ml penicillin, and 100 µg/ml streptomycin at 37 °C. The following day indicated plasmids were transfected using Lipofectamine 2000 (Invitrogen) transfection reagent (2.1 µl Lipofectamine per 1 µg of DNA). Total DNA varied from 700 to 900 ng per well and included EYFP control vector as necessary to maintain equal amount of DNA per well within individual experiments. Approximately 18-24 h after transfection, the culture medium was changed to low-inositol Ham’s F-10 medium (Gibco) containing 1.5 µCi/well Myo-[2-^3^H(N)]inositol (Perkin Elmer) for 16-18h. Accumulation of [^3^H]inositol phosphate was quantitated after the addition of 10 mM LiCl for 1 h to inhibit inositol monophosphate phosphatases. Media were aspirated and cells washed with 1× PBS, followed by the addition of ice-cold 50 mM formic acid to lyse cells. Soluble cell lysates containing [^3^H] inositol phosphate were transferred onto Dowex AGX8 chromatography columns to separate total IP by anion exchange chromatography. Columns were washed with 50 mM formic acid, then 100 mM formic acid and eluted with 1.2 M ammonium formate into scintillation vials containing scintillation fluid and counted. Basal activity is defined as the IP released of the transfected PLC construct relative to the YFP control and in the absence of transfection of any protein activators (G proteins).

### Purification of Gβγ and Gα_q_

Purification of Gβ_1_γ_2_ (Gβγ) was performed by co-expressing Gβγ with His6 Gα_i1_ in High Five insect cells and nickel-agarose chromatography as described previously (Davis et al., 2005; Kozasa and Gilman, 1995). Purification of Gα was performed as described in (Kozasa and Gilman, 1995).

### Purification of His6 PLCβ2, PLCβ2-Δ856, PLCβ2 C-terminal domain

Baculovirus-infected Sf9 cells expressing PLCβ2 and PLCβ2-Δ856, or the C-terminal domain of PLCβ2 were lysed by Dounce on ice in lysis buffer containing 20 mM HEPES, pH 8, 50 mM NaCl, 10 mM β-mercaptoethanol, 0.1 mM EDTA, 0.1 M EGTA, 133 µM PMSF, 21 µg/ml TLCK and TPCK, 0.5 µg/ml Aprotonin, 0.2 µg/ml Leupeptin, 1 µg/ml Pepstatin A, 42 µg/ml TAME, 10 µg/ml SBTI. The lysate was centrifuged at 100,000 × g, and the supernatant was collected and diluted with lysis buffer with NaCl to a final concentration of 800 mM. The diluted supernatant was then centrifuged at 100,000 x g and the supernatant was loaded onto a Ni-NTA column pre-equilibrated with buffer A (20 mM HEPES, pH 8, 100 mM NaCl, 10 mM β-mercaptoethanol, 0.1 mM EDTA, and 0.1 M EGTA). The column was washed with 3 column volumes (CVs) of buffer A, followed by 3 CVs of buffer A supplemented with 300 mM NaCl and 10 mM imidazole. The protein was eluted from the column with 3–10 CVs of buffer A, supplemented with 200 mM imidazole. Proteins were concentrated and loaded onto tandem Superdex 200 columns (10/300 GL; GE Healthcare) equilibrated with SEC buffer (20 mM HEPES, pH 8, 200 mM NaCl, 2 mM DTT, 0.1 mM EDTA, and 0.1 M EGTA). Fractions of interest were confirmed by SDS-PAGE, pooled, concentrated, and flash frozen in liquid nitrogen.

### Biolayer interferometry (BLI)

Protein-Protein interactions were measured by biolayer interferometry using an 8 channel Octet Red system (ForteBio). A 1.5-2 nm shift of PLCβ2-CTD was immobilized on AR2G biosensor tips (Fortebio) by activation with EDC-NHS solution freshly prepared according to manufacturer’s instructions. PLCβ2-Δ856 binding to the immobilized protein was measured at the specified final protein concentrations in buffer containing 20 mM HEPES pH8, 100 mM NaCl, 0.1% BSA and 0.02% Tween, at room temperature and with vigorous shaking (2,200 rpm). Assays were performed at least in triplicate, and binding isotherms were constructed by representing the experimental binding values at equilibrium versus the CTD total concentration.

### Hydrogen deuterium exchange mass spectrometry (HDX-MS)

HDX experiments were performed with or without liposomes (final concentration 0.1 mg/mL) containing brain porcine phosphatidylethanolamine, brain porcine phosphatidylserine and brain porcine phosphatidylinositol 4,5-bisphosphate in a 10:5:1 molar ratio (Avanti Polar Lipids) in a total reaction volume of 50 µl with a final concentration of 400 nM PLC, 1mM NaF, 30µM AlCl_3_ and 600 nM Gα_q_ or Gβγ where appropriate. Deuterium incorporation was initiated by the addition of 40 µl of deuterium oxide buffer (75 mM HEPES pH 7.5, 50 mM NaCl, and 69% (v/v) final deuterium oxide). The constructs were allowed to exchange over three time courses: 3, 30, 300, and 3000s at 16 °C. Exchange was terminated by the addition of 25 µl of ice-cold quench buffer (3% formic acid, 2M guanidine), and samples were immediately frozen in liquid nitrogen.

MS experiments were carried out as described previously (Vadas et al., 2017). In brief, samples were quickly thawed and injected onto an ultra-performance LC system (Dionex Ultimate 3000 RSLCnano system coupled to a leap technologies PAL RTC) at 2 °C. The samples were subjected to two immobilized pepsin columns (Applied Biosystems, Poroszyme, 2-3131-00) at 10 and 2 °C, respectively, at a flow of 200 µl/min for 3 min. Peptides were subsequently collected and desalted on a VanGuard precolumn Trap column (Waters) and were eluted onto an Acquity 1.7 µM particle, 100 × 1 mm2 C18 ultra-performance LC column (Waters). Separation and elution of peptides from the analytical column was achieved using a gradient ranging from 3 to 70% mobile phase B (buffer A: 0.1% formic acid, LC/MS grade; buffer B: 100% acetonitrile, LC/MS grade) over 16 min. Mass spectrometry analyses were performed using an Impact II TOF (Bruker) with an electrospray ionization source operated at a temperature of 200 °C and a spray voltage of 4.5 kV, and data were acquired over a range of 150–2200 m/z. Peptides were identified using the software program PEAKS 7. The false discovery rate was set to 1%. Levels of deuterium incorporation were calculated using the HD-Examiner Software (Sierra Analytics), and data from each individual peptide were inspected for correct charge state, quality of spectra, and the presence of overlapping peptides. Levels of deuteration were computed by HD-Examiner using the centroid of the isotopic cluster corresponding to each peptide at each time point. The deuteration levels calculated are presented as relative levels of deuterium incorporation, the data were further analyzed and curated using Excel, and changes in deuteration levels above 4% and 0.4 Da with a paired t test value of p < 0.05 were deemed to be significant. All deuterium incorporation values for all peptides can be found in the xls file in the source data. The full analysis parameters as required by the IC-HDX-MS (Masson et al., 2019) for all experiments are found in supplemental table 1.

### Liposome Pulldown

200 µM hen egg white phosphatidylethanolamine and 50 µM brain phosphatidylinositol 4,5-bisphosphate (Avanti Polar Lipids) were mixed and dried under N_2_. Lipids were resuspended in sonication buffer (50 mM HEPES, pH 7, 80 mMKCl, 3 mM EGTA, and 2 mM DTT) and sonicated for 2 minutes. Liposomes (130 µL) were combined with 200 ng of PLCβ2 and sonication buffer to a final volume of 200 µL in the presence or absence of 1 μM CTD and/or 500 ng Gβγ. All samples were incubated for 1 h on ice. Samples were then centrifuged at 100,000 × *g* for 1 h. The supernatant was removed, and the pellet was resuspended in 200 µl of sonication buffer. For analysis by SDS-PAGE and Image Studio, 16 µL of the supernatant, or resuspended pellet was denatured with 4 µl of 5× SDS loading dye, and 10 µL of the total sample was analyzed by SDS-PAGE using anti-PLCβ2 antibody (Invitrogen Ca#PA5-75551).

### Phospholipase C activity assay

The assay was performed as described previously (Bonacci et al., 2005), with minor modification. Briefly, lipids containing ∼5000 cpm of [3H-inositol]PIP_2_, 37.5 µM PIP_2_, and 150 µM phosphatidylethanolamine were mixed with 1.4 ng of FL PLCβ2 and equimolar amounts of Δ856-PLCβ2 (1 ng). Varying concentrations of Gβγ were added to the PLC and lipid mixture. The reaction was set at 30 °C for 30 min in the presence of 1.5 mM CaCl2 and 3 mM EGTA (Free Ca^2+^=100nM), and quenched using ice cold 10% TCA. Released soluble [^3^H] IP_3_ was separated through the addition of 5% BSA and centrifugation then measured by liquid scintillation counting. Basal activity is defined as the activity in the presence of Ca^2+^ and no other protein activators.

## Acknowledgements.

Work in the AVS laboratory was supported by a grant from the National Institutes of Health R35GM127303. Work in the JEB laboratory was supported by NSERC through a discovery grant (NSERC-2014-05218) and NSERC Canada Graduate Scholarship-Masters to MLJ.

---

**Supplemental Figure 1:**
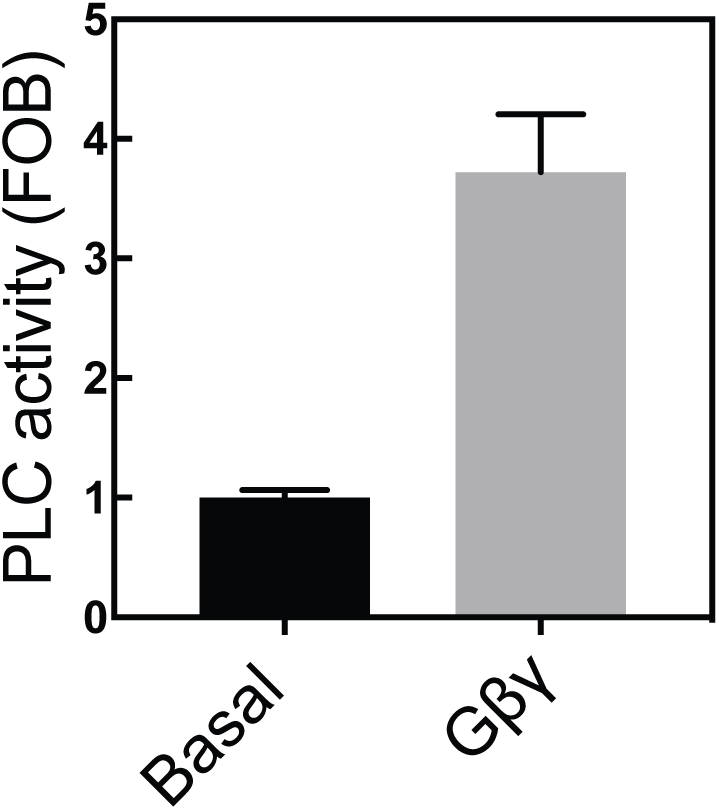
PLCβ2 is activatable by Gβγ under HDX conditions. Reconstituted assay of PLC enzymatic activity as described in in methods section with the exception that reaction was only allowed to proceed for 30 seconds under HDX conditions (200nM PLC, 600nM free Ca^2+^ and 600nM purified Gβγ subunits) Experiments were performed at least in duplicate. The data shown are mean ± S.D

**Supplemental Figure 2:**
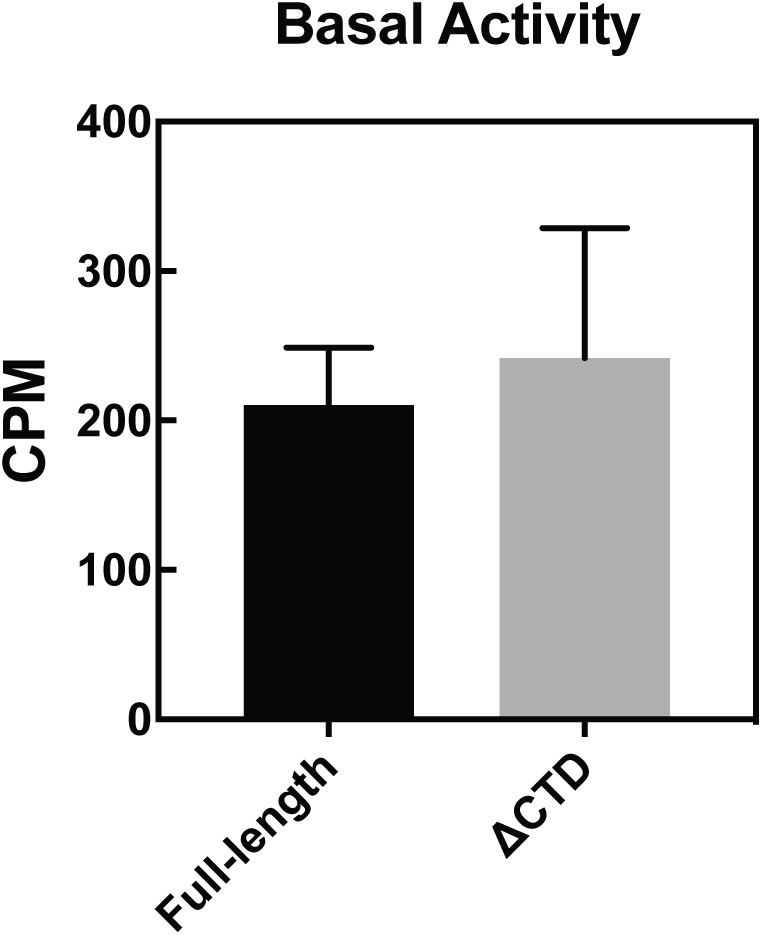
Full length PLCβ2 and PLCβ2-Δ856 have similar basal activities. Reconstituted assay of PLC enzymatic activity unstimulated activity for the PLCβ-Δ856 and Full length PLCβ2 in the absence of any activators. Experiments performed at least 3 times in duplicate. The data shown are mean ± S.E.

**Supplemental Table 1.** Full statistics on all hydrogen deuterium exchange experiments according to the guidelines from the International Conference on HDX-MS.

| Data Set | PLCβ2 | PLCβ2<br>membrane | PLCβ2<br>membrane<br>G$\beta\gamma$ | PLCβ2<br>membrane G$\alpha_q$ |
| --- | --- | --- | --- | --- |
| HDX reaction<br>details | %D<sub>2</sub>O=77%<br>pH<sub>(read)</sub>= 7.5<br>Temp= 16°C | %D<sub>2</sub>O=77%<br>pH<sub>(read)</sub>= 7.5<br>Temp= 16°C | %D<sub>2</sub>O=77%<br>pH<sub>(read)</sub>= 7.5<br>Temp= 16°C | %D<sub>2</sub>O=77%<br>pH<sub>(read)</sub>= 7.5<br>Temp= 16°C |
| HDX time course | 3, 30, 300 and<br>3000 sec at<br>23°C | 3, 30, 300 and<br>3000 sec at<br>23°C | 3, 30, 300 and<br>3000 sec at<br>23°C | 3, 30, 300 and<br>3000 sec at<br>23°C |
| HDX controls | N/A | N/A | N/A | N/A |
| Back-exchange |  |  |  |  |
| Number of<br>peptides | 232 | 232 | 232 | 232 |
| Sequence<br>coverage | 91.6% | 91.6% | 91.6% | 91.6% |
| Average peptide<br>length/<br>redundancy | Length = 12.1<br>Redundancy =<br>4.5 | Length = 12.1<br>Redundancy =<br>4.5 | Length = 12.1<br>Redundancy =<br>4.5 | Length = 12.1<br>Redundancy =<br>4.5 |
| Replicates | 3 | 3 | 3 | 3 |
| Repeatability | Average<br>StDev =<br>0.67% | Average<br>StDev =<br>0.67% | Average<br>StDev =<br>0.67% | Average StDev<br>= 0.67% |
| Significant<br>differences in<br>HDX | >4% and >0.4<br>Da and<br>unpaired t-test<br><0.05 | >4% and >0.4<br>Da and<br>unpaired t-test<br><0.05 | >4% and >0.4<br>Da and<br>unpaired t-test<br><0.05 |  |

